# Revealing the range of maximum likelihood estimates in the admixture model

**DOI:** 10.1101/2024.10.18.619150

**Authors:** Carola Sophia Heinzel, Franz Baumdicker, Peter Pfaffelhuber

## Abstract

Many ancestry inference tools, including STRUCTURE and ADMIXTURE, rely on the admixture model to infer both, allele frequencies *p* and individual admixture proportions *q* for a collection of individuals relative to a set of hypothetical ancestral populations. We show that under realistic conditions the likelihood in the admixture model is typically flat in some direction around a maximum likelihood estimate 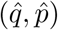. In particular, the maximum likelihood estimator is non-unique and there is a complete spectrum of possible estimates. Common inference tools typically identify only a few points within this spectrum.

We provide an algorithm which computes the set of equally likely 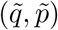, when starting from 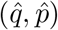. It is analytic for *K* = 2 ancestral populations and numeric for *K >* 2. We apply our algorithm to data from the 1000 genomes project, and show that inter-European estimators of *q* can come with a large set of equally likely possibilities. In general, markers with large allele frequency differences between populations in combination with individuals with concentrated admixture proportions lead to small areas with a flat likelihood.

Our findings imply that care must be taken when interpreting results from STRUCTURE and ADMIXTURE if populations are not separated well enough.

## 1 Introduction

Inferring the ancestry of a sample of individuals from genetic data is a formidable challenge, given its importance in various domains, such as exploration of human history (Rosenberg et al., 2002), identification of missing persons (Phillips, 2015), corrections for population stratification (Pritchard and Donnelly, 2001), forensic genetics (Tvedebrink, 2022) or conservation genetics (Wasser et al., 2007). In these applications, individual genomes are assumed to originate from different ancestral populations with distinct allele frequencies. Both, model-based methods and model-free techniques were developed for estimating individual admixture (Wollstein and Lao, 2015). Model-based methods, such as STRUCTURE (Pritchard et al., 2000), ADMIXTURE (Alexander et al., 2009) and improvements of them (Alexander and Lange, 2011; Shringarpure et al., 2016; Dominguez Mantes et al., 2023; Falush et al., 2003, 2007; Hubisz et al., 2009; Raj et al., 2014; Ko et al., 2023) treat the number of ancestral populations (usually denoted by *K*), allele frequencies within populations and individual admixture of all sample individuals as parameters of a statistical model, which we will call the admixture model. Model-free methods use for example Principal Component Analysis or Spectral Graph Theory (Klei et al., 2011; Lee et al., 2010; Jombart et al., 2010) for inferring the individual ancestry (see Wollstein and Lao (2015) for an overview) and links between these two approaches have been made (Engelhardt and Stephens, 2010). In recent years, neural networks have become yet another tool for inference of individual admixture, which follow the admixture model (Dominguez Mantes et al., 2023), or similar research questions (Battey et al., 2020).

There are various cases where the output of STRUCTURE and ADMIXTURE must be carefully interpreted. Different historic scenarios might lead to similar genetic data, and researchers have been warned not to over interpret estimates of the admixture model (Lawson et al., 2018). In addition, population structure is not correctly recovered in all cases (Puechmaille, 2016) and it needs to be evaluated whether the assumptions of the admixture model are met, Garcia-Erill and Albrechtsen (2020). Another discussion in the admixture model comes from the choice of *K*. Since the *K* populations are ancestral, it is per se unclear what the right choice of *K* is, given an unstructured sample of genetic material (Evanno et al., 2005; Pritchard et al., 2000). As the admixture model does not account for the potential relationships between ancestral populations, it can result in contradictory predictions when using different values for *K* (Burger et al., 2024). *Frequently, inferring K* from the data leads to the choice *K* = 2, see e.g. (Janes et al., 2017; Friedlaender et al., 2008; Rosenberg et al., 2005; Lawson et al., 2018; Hamlin and Arnold, 2014; Phillips et al., 2009), while it is questioned if *K* can reasonably inferred at all (Verity and Nichols, 2016; Wang, 2017). Due to the increasing number of parameters, STRUCTURE is in practice not applied for many different populations, i.e. *K* ≥ 10 (Lawson et al., 2012).

In the present study, we focus on principal limitations of the admixture model from a statistical perspective. Usually, STRUCTURE and ADMIXTURE result in maximum likelihood estimators (or maximum aposteriori estimators) of both, allele frequencies and admixture proportions. In a sample of *N* individuals, *M* bi-allelic markers and *K* ancestral populations, we have to estimate both, the *N* × *K*-matrix of individual admixtures, and the *K* × *M* -matrix of allele frequencies.

One of the well-known limitation of the admixture model is usually called <u>label-switching</u>, which means that relabeling ancestral populations leads to equivalent results. This corresponds to applying the same permutation to the population indices in both matrices. Label-switching is of practical importance, since it is one reason why different runs of STRUCTURE or ADMIXTURE can result in different estimates (Jakobsson and Rosenberg, 2007; Alexander et al., 2009; Pritchard et al., 2010; Verdu et al., 2014; Kopelman et al., 2015; Behr et al., 2016; Cabreros and Storey, 2019; van Waaij, 2022; Liu et al., 2024). As a possible solution, a series of investigations tries to resolve label-switching with high computational effort in order to be able to compare different outputs of STRUCTURE (Behr et al., 2016; Rosenberg, 2004; Jakobsson and Rosenberg, 2007; Earl and VonHoldt, 2012; Kopelman et al., 2015), even for different choices of *K* (Margaryan et al., 2020; *Larena et al*., *2021; Jia et al*., *2020; Liu et al*., *2024)*.

We show that the non-uniqueness of point estimates substantially extends beyond the phenomenon of mere label-switching and can be more severe. The differences between equally likely estimates are not just nuances, they can be substantial. However, the set of equally likely estimators is small if markers are able to distinguish clearly between populations. Therefore, our results highlight that reliable ancestry estimates depend on at least a few SNPs with large allele frequency differences between the populations. If such SNPs are unavailable, estimates within the admixture model must be taken with caution.

The structure of the paper is as follows: After introducing the admixture model, we start our investigation with a single estimator of individual admixtures and allele frequencies, and look for the range of estimators which are equally likely. From this we can compute the differences between all the identified estimators with respect to the estimated allele frequencies and the estimated individual admixtures. This results in an algorithm, EMALAM (Every MAximum Likelihood estimator in the Admixture Model) available at our GitHub repository or as online version that explores the range of equally likely estimators. To show the consequences of the flatness of the likelihood curve, we apply our theoretical results to genetic data from the 1000 Genomes Project (Consortium et al., 2015). Furthermore, we compare our approach to Pong, a method proposed by Behr et al. (2016), i.e. to running STRUCTURE several times while accounting for label-switching to identify different modes. In contrast to the identification of the different modes discovered by STRUCTURE/ADMIXTURE, here we explore the complete space of equally likely estimators.

## 2 Methods

### 2.1 The Admixture Model

*Let us reintroduce the admixture model in the unsupervised setting. For M* markers, where marker *m* has *J*_*m*_ different alleles, *m* = 1, …, *M*, the genetic data of *N* diploid individuals is given by 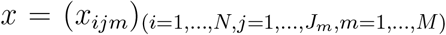. Additionally, *x*_*ijm*_ ∈ {0, 1, 2} determines the number of copies of allele *j* in individual *i* at locus *m*. We assume that we are dealing with diploid individuals, i.e.,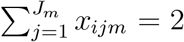. We fix the number of distinct ancestral populations *K* and aim to infer the individual admixtures *q* = (*q*_*ik*_)_*i*=1,…,*N,k*=1,…,*K*_ and the allele frequencies *p* = (*p*_*kjm*_)_*k*=1,…,*K,j*=1,…,*J*_*m,m*=1,…,*M* from the genetic data. Here, *q*_*ik*_ represents the proportion of the genome of individual *i* that originates from population *k* (i.e., the individual ancestry of individual *i* in population *k*), and *p*_*kjm*_ represents the frequencies of allele *j* in the assumed ancestral population *k* at marker *m*.

To depict the likelihood, we write *q*_*i*,·_ for the row vector (*q*_*i*1_, …, *q*_*iK*_), and the allele frequencies as *p*_·*jm*_ = (*p*_1*jm*_, …, *p*_*Kjm*_) and note that *q, p* satisfy the conditions (where 1_*K*_ is the 1-vector of length *K*)

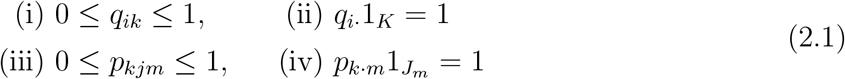

i.e., the individual admixtures *q* of each individual sum to one and allele frequencies *p* sum to one at each locus. Given *x*_*ijm*_, the number of alleles *j* of individual *i* at marker *m*, there is a constant *C*_*x*_ which depends only on *x* = (*x*_*ijm*_), such that the log-likelihood for *p, q* given *x* is, (using ⊤ to indicate column vectors)

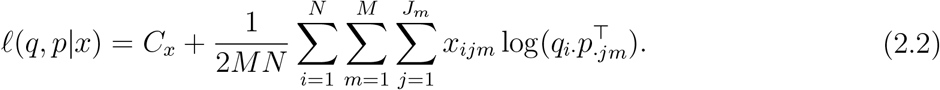

Here, 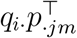 is a scalar product, i.e.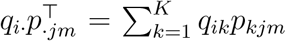. Apparently, the likelihood (2.2) depends on *q, p* only via 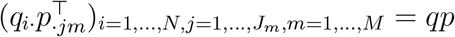. Maximum likelihood estimators are then given by

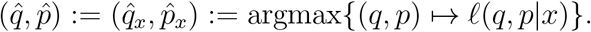

### 2.2 Equally likely estimators

We start with an explanation why there are different estimators in the admixture model with the same likelihood. As stated above, the likelihood *𝓁*(*q, p*|*x*) depends on *q, p* only via the scalar products 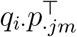. As a consequence, 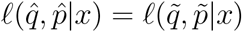 provided that

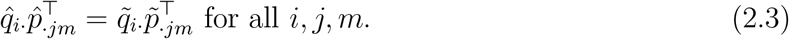

In order to see that this is in fact possible, let *S* be an invertible *K* × *K* matrix as well as

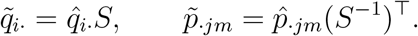

Writing out the scalar product, 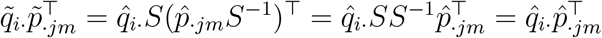, i.e(2.3) holds. However, we have to be careful in order not to violate the side conditions (2.1). In order to achieve this, restrict *S* to a matrix with rows summing to 1, i.e.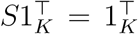. (If *S* has non-negative entries, it is called a stochastic matrix.) Multiplying this equation by *S*^−1^, it is clear that 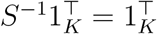 as well. Then, provided 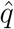 and 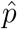 satisfy (2.1), we have

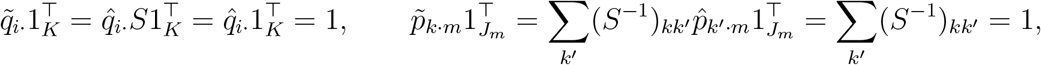

i.e. 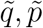 satisfy the side conditions (ii), (iv) as well. Moreover, if *S* is from

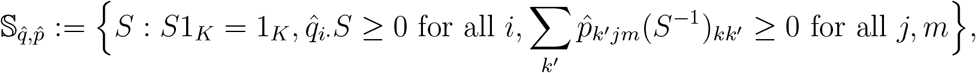

side conditions (i) and (iii) are satisfied for 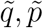 as well. So, we have shown that the likelihood curve is flat in some directions of the parameter space, and hence likelihood-based estimators are not unique provided that 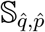 is not trivial. Moreover, we can search 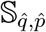 for parameters which lead to equally likely individual admixtures and allele frequencies.

A well-known but not exclusive example of this non-uniqueness deals with label-switching for populations. For this, consider a permutation matrix *S*. (Such a matrix has entries in {0, 1} with a single 1 in each row/column.) Here, *Sq*^⊤^ are individual admixtures where we have switched labels *k* → *l* if (*S*)_*kl*_ = 1. Similarly, labels of allele frequencies *p*, switch label accordingly, i.e. using *S*^−1^*p*^⊤^. For such matrices, conditions (i), (iii) are always met and it has been discussed frequently (see e.g. Behr et al., 2016; Ko et al., 2023) that the estimators *Sq, S*^−1^*p* and *q, p* are equally likely. However, label switching is only one example, and the goal here is to identify the complete range of equally likely estimators beyond label switching.

### 2.3 EMALAM

Our algorithm for exploring the set of equally likely estimators starts with a single estimator 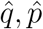, as e.g. provided by the output of STRUCTURE or ADMIXTURE. We then search for 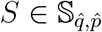 as follows: Pick a subset ℐ ⊆ {1, …, *N*} of individuals and (recalling that *H*(*x*) = − ∑ _*k*_ *x*_*k*_ log *x*_*k*_ is the entropy of *x* which satisfies *x*1^⊤^ = 1, which is maximal for the uniform distribution, and minimal for point measures) either

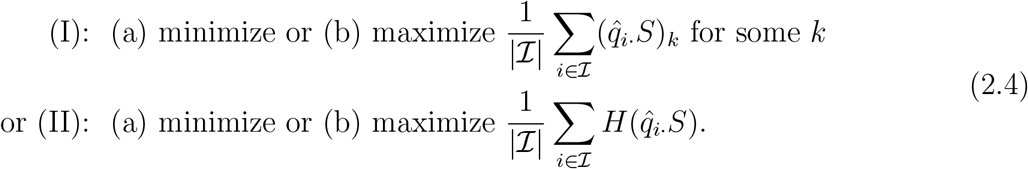

So, in (I), we look at a concrete population *k* and minimize / maximize the average individual ancestry in *k* for all individuals in ℐ. In (II), we rather minimize (favoring less admixed individuals) or maximize (favoring admixed individuals) the average entropy for individuals in ℐ. See also Figure 4 for an illustration how an equally likely estimator looks like for the four different options in an example data set.

### 2.4 Non-identifiability and allele frequency differences

Let us consider a toy example for the claimed non-identifiability. We simulate 1000 markers with different allele frequencies among 100 admixed individuals from two populations. The true individual admixtures are depicted as the black line in Figure 1, and the range of equally likely estimators as given by (3.1) are the colored area. Depending on whether we only have 1000 markers with intermediate allele frequency differences, or additionally two markers with a large frequency differential, the range of equally likely estimators varies; see Figure 1. Recall that markers which are (almost) fixed in one, and polymorphic in all other populations, are called anchor markers. Equivalently, individuals with individual ancestry almost exclusively in one population, are called anchor individuals (Cabreros and Storey, 2019). Figure 1 thus also illustrates how the addition of anchor markers and individuals influences the range of equally likely estimators. Without such anchors, the range of equally likely admixture proportions can span a substantial area such that the contribution of the two populations is basically impossible to disentangle (Figure 1 top left). In the case with two populations A and B, if for an equally likely estimator the admixture proportion for population A of a certain individual is increased, so is the admixture proportion for population A in all other individuals, and the allele frequencies change accordingly. However, when *K >* 2 the dependencies will be more complex.

**Figure 1.**
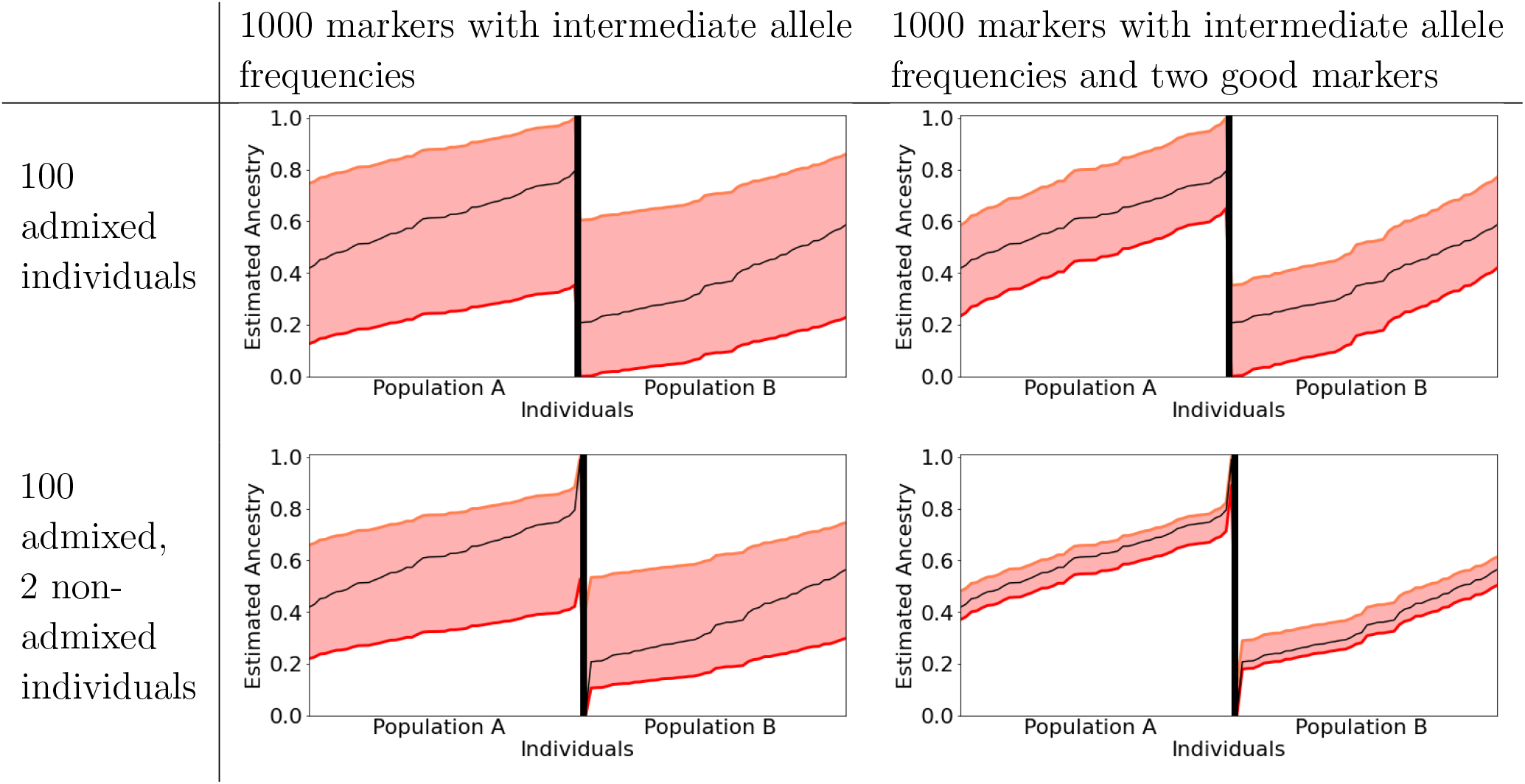
Range of equally likely estimators for two simulated populations. The *x*-axis represents the individuals and the *y*-axis indicates the individual ancestry for population A. The black lines show the true (simulated) individual ancestries. The area between the red lines symbolizes equally likely estimators. The estimated allele frequencies in population A for 1000 bi-allelic markers were simulated according to a uniform distribution in [0.25, 0.75]. The allele frequencies in population B are *p*_1,1,*m*_ + *ϵ*, where *ϵ* is uniformly distributed in [0, 0.3] if *p*_2,1,*m*_ ∈ [0.2, 0.8] and *p*_2,1,*m*_ = 0.5 else. Additionally, we added in the right column two additional markers with frequencies (0.01, 0.1), (0.9, 0.99) in populations A and B, respectively. In the second row, we included two non-admixed individuals with ancestry (0.01, 0.99) and (0.99, 0.01).

### 2.5 Data applications

Since our main focus is on a proof of concept of non-identifiability in the admixture model, we use genetic data from the 1000 genomes project (Consortium et al., 2015). In our applications, we choose every 10.000^*th*^ bi-allelic SNP among all SNPs with a minor allele frequency of 5%, resulting in about 10^5^ markers for intercontinental and about 7 · 10^4^ for intra-continental applications. Moreover, we considered the Kidd AIM set (Kidd et al., 2014) with 55 bi-allelic markers.

## 3 Results

### 3.1 Some theoretical insights

**The case** *K* = 2

The case *K* = 2 is important from an application perspective (Friedlaender et al., 2008; Rosenberg et al., 2005; Lawson et al., 2018; Hamlin and Arnold, 2014; Phillips et al., 2009). In this case, starting with an estimator 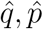, we give for each individual *i* the range of possible individual admixtures with the same likelihood.

We show in Theorem 1 in Appendix A that equally likely estimators for individual ancestry of individual *i* in population 1, i.e. *q*_*i*1_, are in

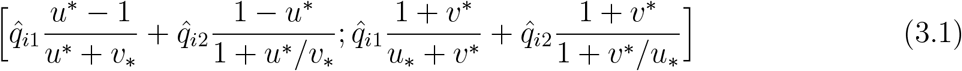

with

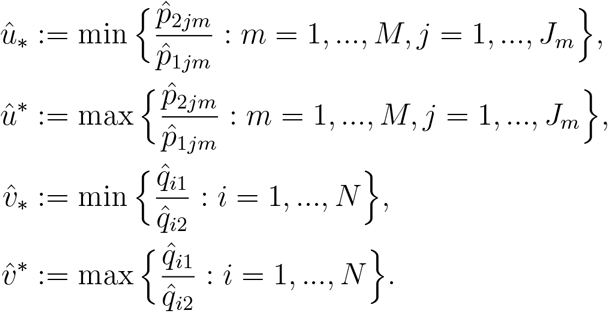

Most notably, the range of possible admixture proportions of individual *i* among the equally likely estimators only depends on 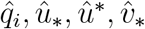 and 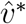. So, adding markers *m*^′^ or individuals *i*^′^ with less extreme quotients 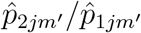 and 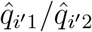 than markers/individuals already present in the dataset does not solve the problem of non-identifiability per se. From the formula, the lower bound is near 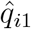 provided that both, *v*_∗_ ≪ 1 ≪ *u*^∗^, and the upper bound is close to 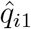 iff *u*_∗_ ≪ 1 ≪ *v*^∗^. In other words, we need both, markers which can distinguish well between both populations, leading to *u*_∗_ ≪ 1 ≪ *u*^∗^, as well as individuals which are almost non-admixed for both populations, leading to *v*_∗_ ≪ 1 ≪ *v*^∗^. This gives a more quantitative view on the fact that anchor individuals and anchor markers are necessary for uniqueness of the equally likely estimator; see the anchor conditions from Cabreros and Storey (2019).

**The case** *K >* 2

In this case, we did not obtain an analytical solution for the possible individual admixtures. However, we note that only changing 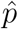 and 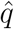 in two populations could be treated in the same way as the case *K* = 2. However, this only leads to a lower bound for the set of possible individual admixtures. Therefore, we rely on numerical minimizations and maximizations in cases (I) and (II) for *K >* 2 in order to find extreme, but equally likely estimators 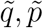. Some more information can be found in the Documentation of EMALAM.

### 3.2 Non-identifiability in the 1000 genomes data

We apply EMALAM to the data from the 1000 Genomes Project (Consortium et al., 2015). Additionally, we consider the difference between our method and running STRUCTURE many times as proposed by Behr et al. (2016).

In our workflow, we run STRUCTURE, and then study the range of equally likely estimators using the STRUCTURE estimator and EMALAM. We use data from northern Europeans from Utah (CEU, 99 individuals), Great Britain (GBR, 91 individuals), Iberian Populations in Spain (IBS, 107 individuals) and Tuscans from Italy (TSI, 107 individuals) in different combinations in the following. For the (small) marker set from Kidd et al. (2014), we make 20 independent runs of STRUCTURE, whereas we make only one run for the large marker set.

First, we consider two ancestral populations, *K* = 2. Using only data from GBR and IBS, Figures 2 and 3 show the range of equally likely individual admixtures (the colored area). In order to put our results into context, we (i) run STRUCTURE 20 times and apply pong (Behr et al., 2016) to the outputs to avoid label switching for the small marker set and (ii) compare the area of equally likely estimators when starting with the outputs of STRUCTURE and ADMIXTURE for the larger marker set. For (i), consider the blue and green lines in Figure 2, which uses the marker set from Kidd et al. (2014). Here, the differences between the 20 runs of STRUCTURE are much smaller than the differences between the equally likely estimators found by EMALAM.

**Figure 2.**
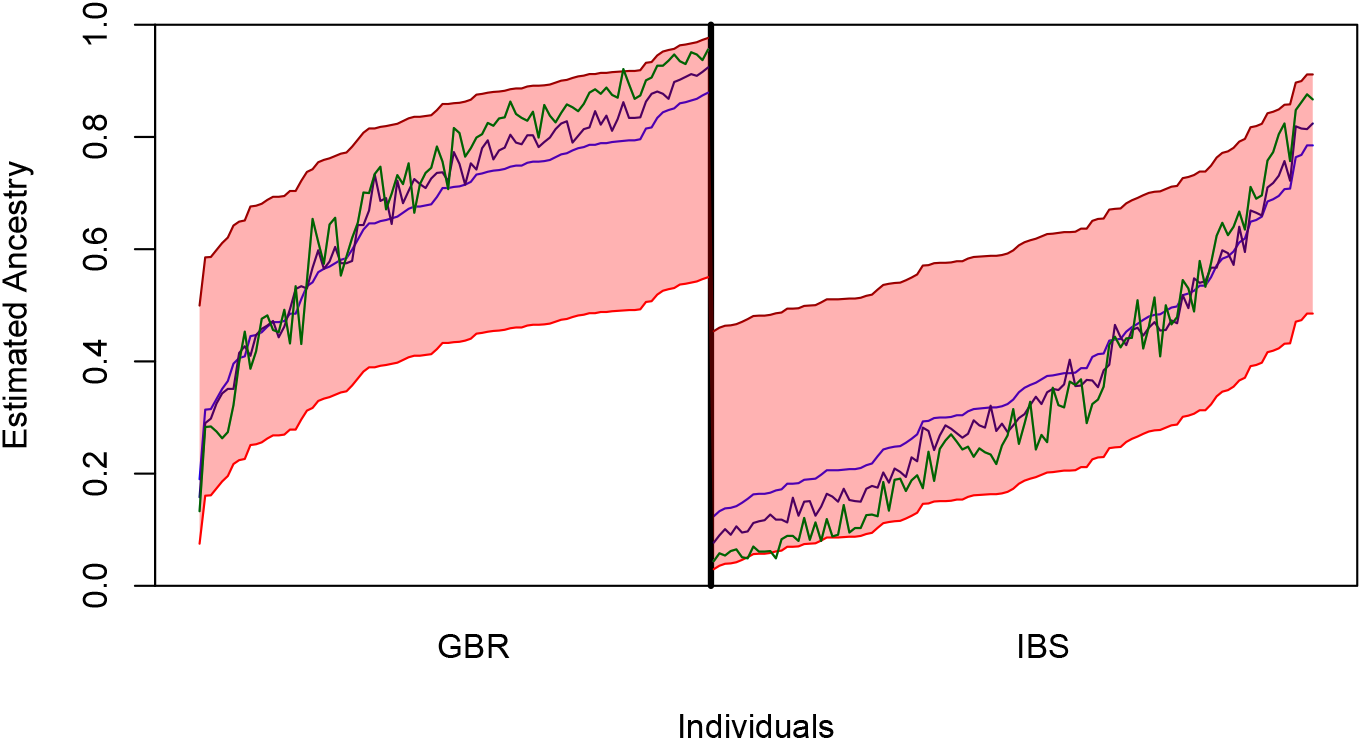
Range of equally likely estimators for GBR and IBS in the admixture model with *K* = 2 with the marker set from Kidd et al. (2014). The colored area is between the minimal and maximal equally likely estimator for the individual ancestry, if we apply the formula (3.1). The three colored lines depict the output of 20 runs of STRUCTURE. The software pong was used to only depict the three result modes that are not similar. Here, the green and dark blue line represent minor modes, while the blue line represents the major mode.

**Figure 3.**
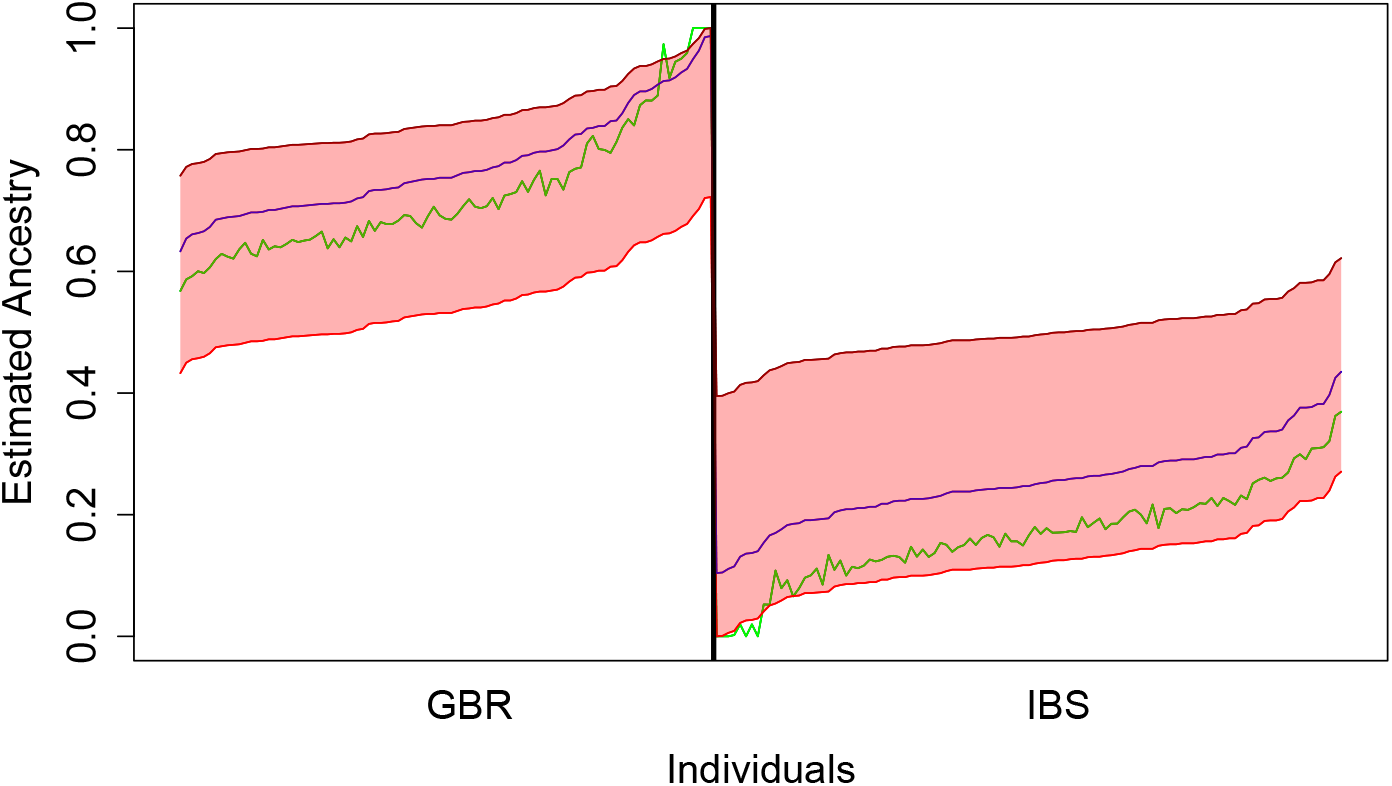
Range of equally likely estimators for GBR and IBS populations in the admixture model with *K* = 2. Ancestry proportions of the estimators are depicted as in Figure 2 but here we use 71185 SNPs. We only consider bi-allelic SNPs, where the allele frequency of both alleles is higher than 0.05. Additionally, we used only every 10000^*th*^ SNP. The green line corresponds to the estimated ancestry of ADMIXTURE. The blue line is the output of STRUCTURE.

**Figure 4.**
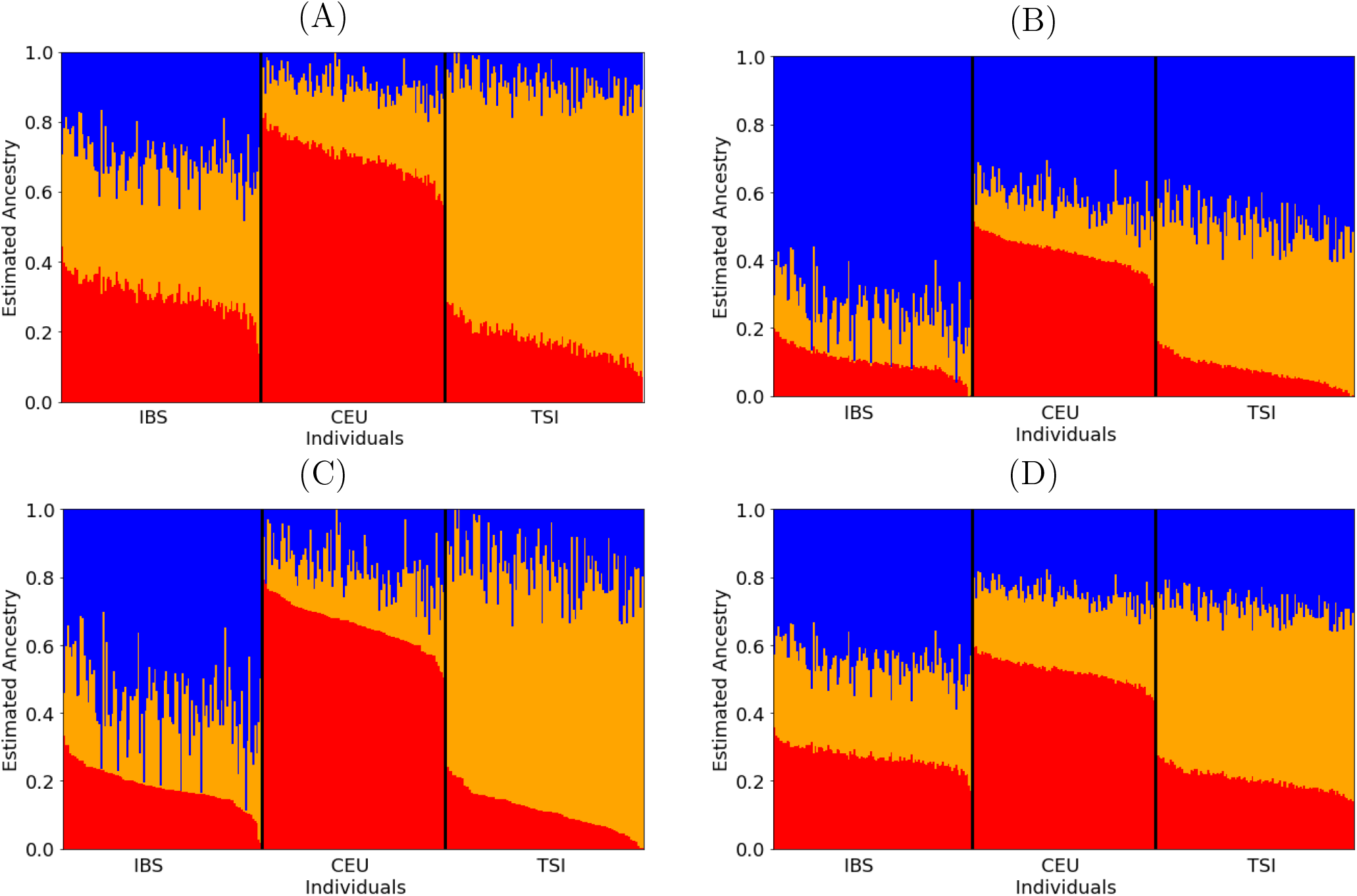
Estimated individual admixtures for the TSI, IBS, and CEU populations with 125350 markers. All four figures show estimators with the same likelihood. We used EMALAM with *K* = 3 in order to optimize the following target in the four figures: (A) Minimize the ancestry from the blue population (option (I)(a)); (B) Maximize the ancestry from the blue population (option (I)(b)); (C) Minimize the entropy (favoring non-admixed individuals), i.e. option (II)(a); (D) Maximize the entropy (favoring admixed individuals), i.e. option (II)(b). Orders of individuals on the x-axis is the same for (A)–(D).

For (ii) consider the same individuals in Figure 3, but using the large marker set of 71185 SNPs. Here we see that the range of estimators with the same likelihood as the output of STRUCTURE is still large, comparable with the range when using the marker set from Kidd et al. (2014) (Figure 2). We also see that the difference between the outputs of STRUCTURE and ADMIXTURE is small. Interestingly, we see that the difference between the maximal and the minimal estimator with the same likelihood as the output of ADMIXTURE is much smaller, i.e. so small that we cannot see any differences in Figure 2.

Second, we consider another combination of populations, namely CEU, TSI and IBS where we apply EMALAM to the data with *K* = 3. In Figure 4, the data was created by running STRUCTURE once and by applying option (I) (a) and (b) (recall from (2.4)) to the blue population and option (II) (a) and (b) to the output of STRUCTURE for *K* = 3. The different options of EMALAM aim to highlight different aspects where the interpretation of STRUCTURE and ADMIXTURE results can be particularly misleading when the complete range of estimators with equal likelihood is not considered. Option (I) maximizes and minimizes the ancestry proportions of a particular population, which can be useful to assess the confidence with which an individual is attributed to a certain bio-geographic background. Option (II) can highlight a different aspect as it maximizes or minimizes the entropy of admixtures in all individuals. This option thus helps to assess the identifiability of separate populations and whether the individuals can be separated into multiple populations. This option helps determine if the estimators clearly suggest that individuals can be categorized into distinct populations or whether individuals are more admixed.

## 4 Discussion

We demonstrated how the non-uniqueness of the estimators in the admixture model impacts the range of equally likely estimators.

Here, we introduced EMALAM, a method to explore the complete range of equally likely estimators. In addition, EMALAM is able to merge estimators that occur due to label switching; cf. pong (Behr et al., 2016). We stress that in general, the non-uniqueness of equally likely estimators has way more possibilities than just switching labels. Since EMALAM relies on a numerical minimization of a *K* × *K*-matrix, it is faster than running STRUCTURE or ADMIXTURE multiple times. Furthermore, it identifies the most extreme estimators with respect to specific aspects, i.e. the individual admixtures of a single individual or the admixture of a complete population.

In general, anchor markers, which have fixed alleles in only one population, and anchor individuals with non-admixed ancestry, play a vital role in identifiability of the admixture model. If such anchors are missing, the range of estimators which EMALAM computes can be too large for a useful interpretation of the results provided by STRUCTURE or ADMIXTURE concerning the inference of the population structure, such that for some MLEs the ancestry fractions of one populations can vanish. Frequently, the lack of anchors goes hand in hand with choosing a too large number of ancestral populations. It is easy to see that the number of optima increases if *K* increases, i.e. in the admixture model *K* should be chosen as small as possible to prevent non-uniqueness. However, if suitable markers and individuals and a small enough *K* are considered the difference between equally likely estimators can be small (see Figure 1).

EMALAM helps to investigate the range of equally likely estimators and how meaningful the inferred ancestries are. As a rule of thumb, small differences between the estimated individual ancestries and the estimated allele frequencies lead to a large range of possible estimated individual admixtures with the same likelihood as the output of STRUCTURE.

Our results highlight that a larger number of markers alone does not necessarily lead to a meaningful individual ancestry inference in the admixture model. It is crucial to include at least some markers with high allele differences between the populations of interest and to include for each population individuals with low admixture proportions. This implies that even when using thousands of markers (or individuals), the certainty of individual ancestry inference in the admixture model can rely on only a few ancestry informative markers (Figure 1). Consequently, for populations where admixture occurs frequently or no informative markers exist the range of equally likely estimators in the admixture model can be larger than considering the different modes of multiple STRUCTURE runs suggests.

One caveat of applying EMALAM is that it only identifies equally likely estimators starting from a given initial estimator. Estimators with similar or even lower likelihoods are not detected, and the resulting range of equally likely estimators depends thus on the starting point. For example, if an estimator inferred by ADMIXTURE creates a spurious anchor individual or allele, the inferred range may appear narrow, which could potentially give a false impression of high confidence. While this is an extreme case, it is advisable to run STRUCTURE/ADMIXTURE multiple times to capture the variability in inferred modes, and apply EMALAM to each mode to better assess the range of equally likely estimators for each of them.

Using alternative models and methods that restrict the parameter space, e.g., by considering linkage disequilibrium and local ancestry (Falush et al., 2003; Corbett-Detig and Nielsen, 2017), or by assuming dependency structures between the ancestral populations (Bradburd et al., 2018; Burger et al., 2024) can circumvent the non-identifiability of the admixture model. However, while more complex models can lead to more precise individual admixtures and allele frequency estimates, they typically rely on a larger number of markers or specific model assumptions. Instead, EMALAM can be used to assess the consequences of the non-identifiable equally likely estimators in applications based on the admixture model. Especially for scenarios with small allele frequency differences between populations EMALAM will help to prevent misinterpretation of ADMIXTURE and STRUCTURE results.

## 6 Data Availability Statement

The data is available at the 1000 Genomes Project website (Consortium et al., 2015). The implementation of EMALAM can be downloaded from the GitHub repository. An online version is also available.

## 7 Acknowledgments

We thank Jakob Stiefel for some early work on this project.

## Supplemental Material

### A Theoretical results

#### A.1 Two ancestral populations

Of course, *K* = 2 in this section, so we suppress indices of 2 wherever possible. In addition, we assume we are given some 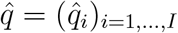, i.e. some estimator for all individual admixtures for all individuals, as well as 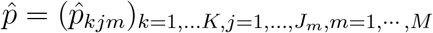, i.e. all estimated allele frequencies at all markers satisfying (2.1). We will use vector notation whenever possible, e.g. *p* ≥ 0 for some vector *p* means that all entries in *p* are ≥ 0. We start by noting that for 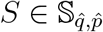, we will require 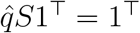, which we achieve by assuming that *S*1^⊤^ = 1^⊤^, so we will work with *S* and the corresponding *S*^−1^ of the form

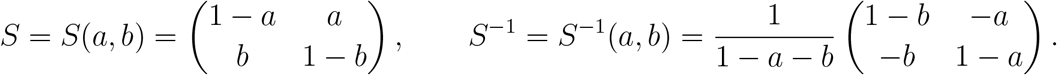

For such *S*, we set

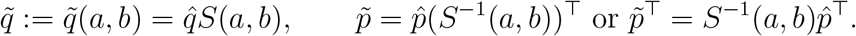

Since 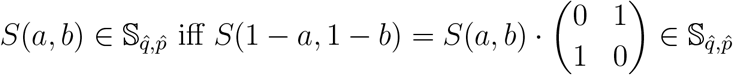 (i.e. switching population labels leads to an additional solution), we prevent finding the solution with switched labels by using the condition *a* + *b* ≤ 1, i.e. we do not consider the additional solution (1 − *a*, 1 − *b*). So, our goal is to determine the set

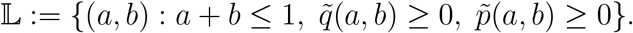

(Since summing *q* (over *k*) and *p* (over *j*) equals one as explained in Section 2, this suffices also for 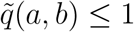 and 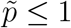.)

For the statement of our Theorem, we need the following notation:

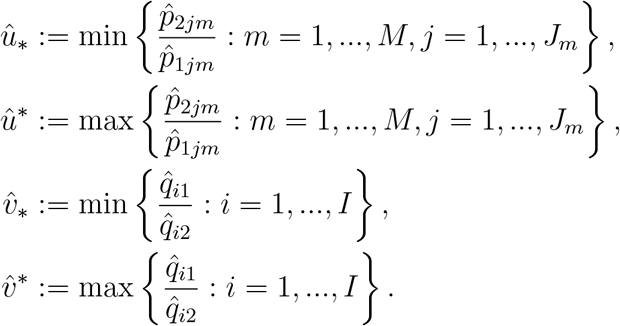

##### Theorem 1

(Range for 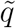 and 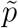). *The set* 𝕃 *is as given in Figure S1*. *Moreover*, (*a*^∗^, *b*^∗^) *and* (*a*_∗_, *b*_∗_) *maximizing and minimizing* 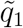 *in all individuals are*

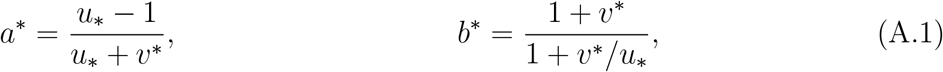

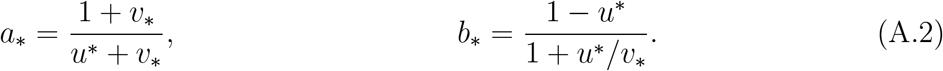

*and for individual i, the resulting possible* 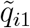 *are*

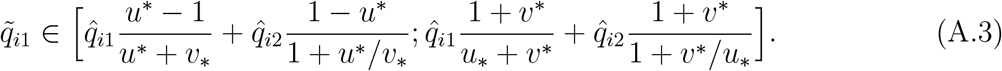

*In addition, the resulting possible* 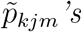 *are*

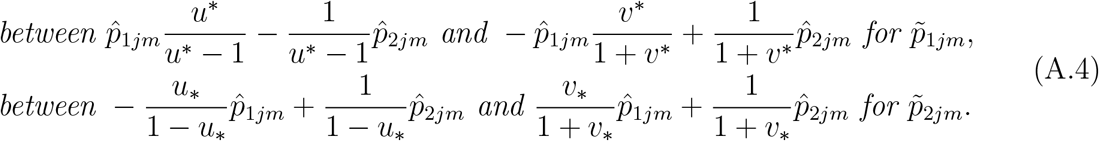

*Proof*.

**Step 0: Computing** 𝕃: As explained in Section 2, our task is to find *S* such that 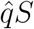 and 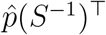 are non-negative. So, for *S* = *S*(*a, b*), we find for 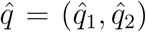 and 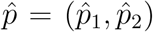 the conditions

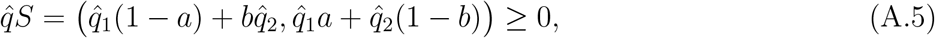

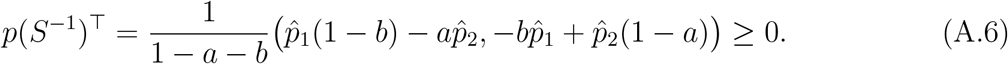

From these equations, we find the equivalent conditions (note that we are using *a* + *b* ≤ 1 in the second line)

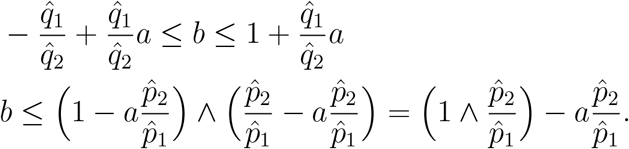

Since all conditions are linear, they are satisfied if and only if they hold for the maximal and minimal possible values for 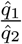 and 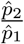, i.e.

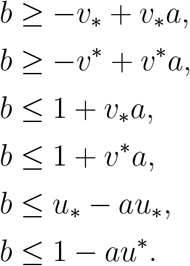

We have used that 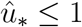 and 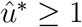 by construction: if 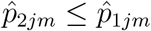 for some *j, m*, there must be *j*^′^ with 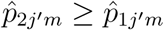 since summing over *j* is 1. We have illustrated all inequalities and the resulting L in Figure S1. Note that *a* + *b* ≤ 1 is satisfied in 𝕃.

**Step 1: Maximizing and minimizing** 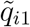: We have to find combinations (*a, b*) ∈ 𝕃 minimizing and maximizing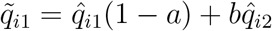.

In order to maximize 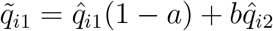, note that a maximizer (*a*^∗^, *b*^∗^) ∈ 𝕃 must have the property that no (*a*^′^, *b*^′^) ∈ 𝕃 with *a*^′^ *< a*^∗^, *b*^′^ *> b* exists (since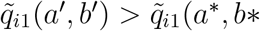) in this case, contradicting the maximization property of (*a*^∗^, *b*^∗^)). This already implies that (*a*^∗^, *b*^∗^) is located on the line *b* = 1 + *v*^∗^*a*, since it is the line bounding 𝕃 in the north-west. We find along this line

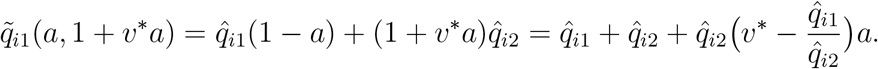

Since 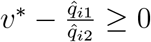 by construction, 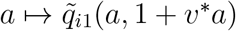 is increasing and the maximizer is at

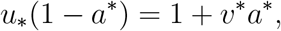

i.e.

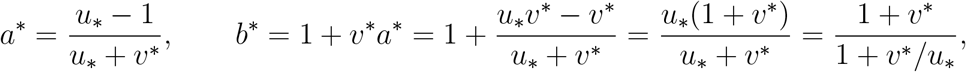

which shows (A.1).

For the minimizer of 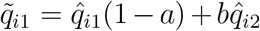, we analogously note that a minimizer (*a*_∗_, *b*_∗_) ∈ 𝕃 must have the property that there is no (*a*^′^, *b*^′^) ∈ 𝕃 with *a*^′^ *> a*^∗^, *b*^′^ *< b*. This implies that (*a*_∗_, *b*_∗_) is located on *b* = −*v*_∗_ + *v*_∗_*a*, since this is the south-eastern boundary of 𝕃. We find

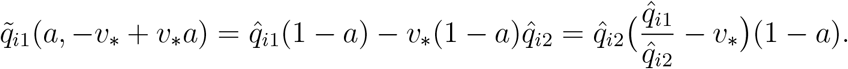

Since 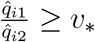 by construction, 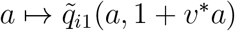 is decreasing and the minimizer (*a*_∗_, *b*_∗_) is at

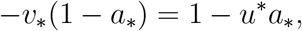

i.e.

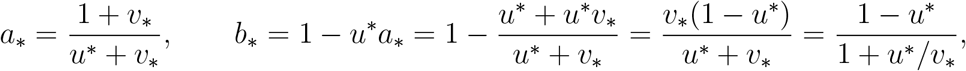

giving (A.2).

Plugging (*a*^∗^, *b*^∗^) into 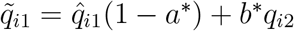, we obtain the maximal 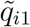, which is given by

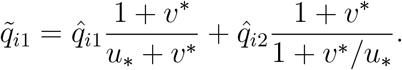

Accordingly, we obtain the minimal 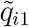 as

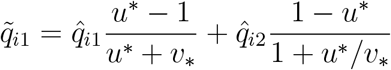

This gives (A.3) and we have shown all claims for the range of 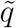.

**Step 2: Maximizing and minimizing** 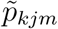: Here, we have to find combinations (*a, b*) ∈ 𝕃 minimizing and maximizing 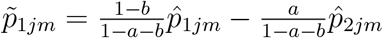 and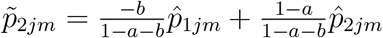. We start by a reparametrization of *S* with

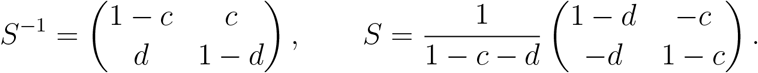

Since we restrict to *a* + *b* ≤ 1, we must assume

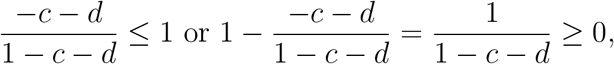

which implies *c* + *d* ≤ 1. So, we impose the restriction *c* + *d* ≤ 1 in the sequel. In other words, we set

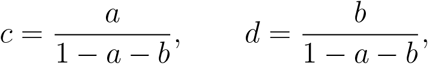

or

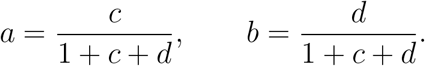

The conditions (A.5) and (A.6) become

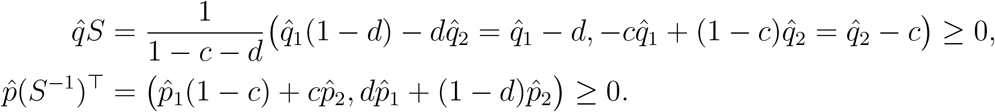

Since *c* + *d* ≤ 1, we have

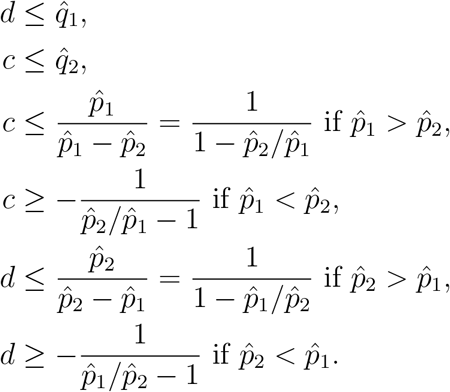

The first three inequalities have to hold for all *i*, so we can write

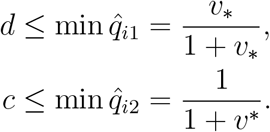

The latter four inequalities have to hold for all *j, m*, we can replace them by

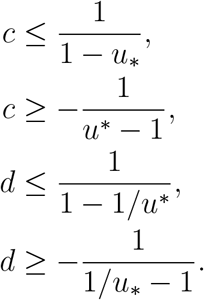

An illustration of all points (*c, d*) satisfying these inequalities is given in FigureS2.

In order to maximize (or minimize) 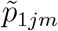, we have to distinguish two cases. If 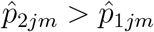, we have to find the maximal (minimal) allowed *c*. If 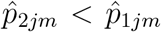, we must rather find the minimal (maximal) allowed *c*. In each case, 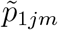 is between

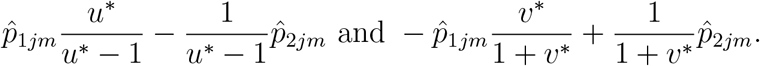

Similarly, 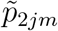 is between

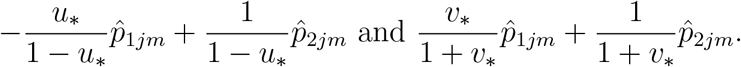

This gives (A.4) and we have shown all claims.

**Figure S1:**
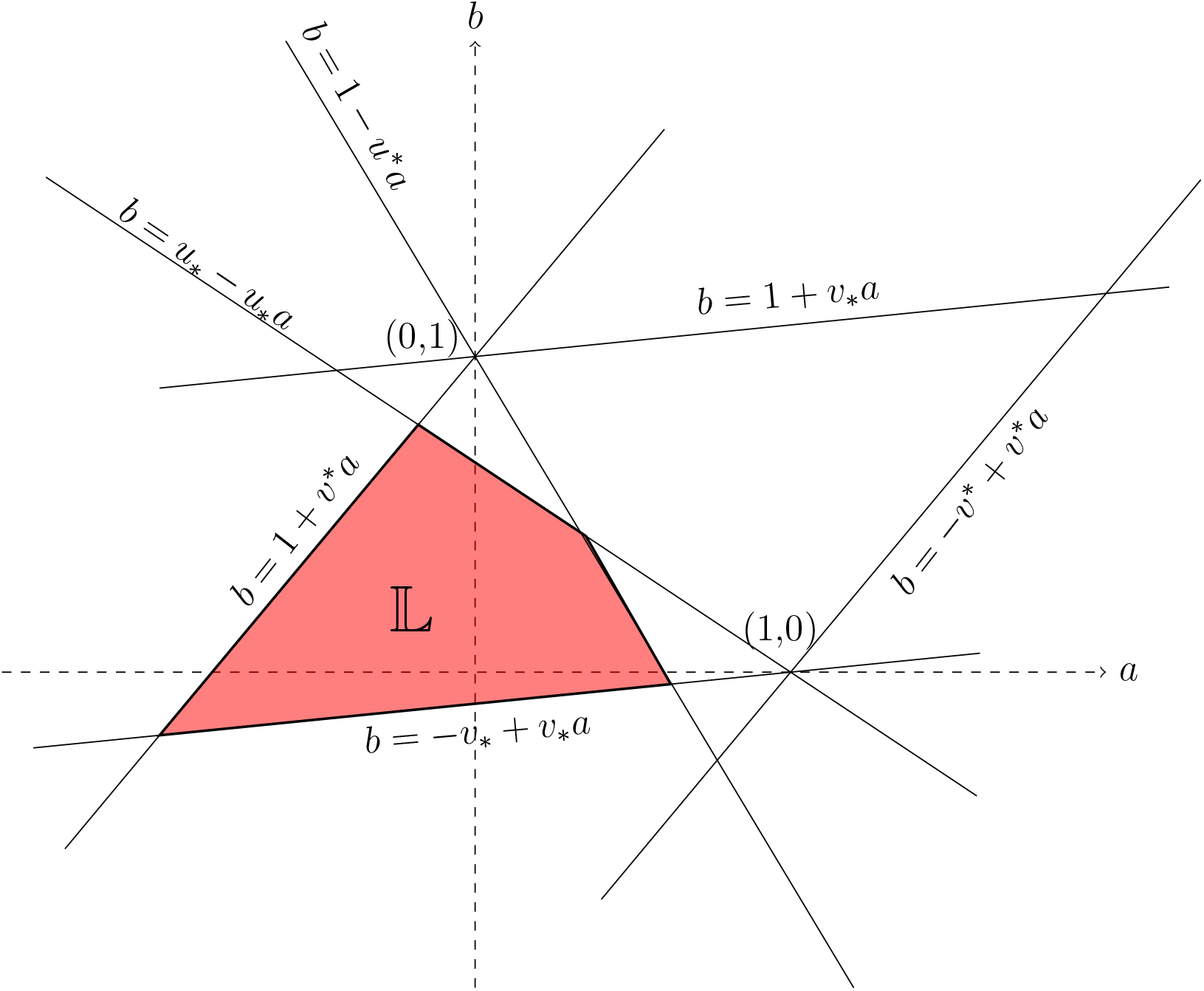
The set of (*a, b*), such that 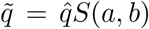 and 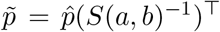 satisfies the conditions (2.1), is given by 𝕃.

**Figure S2:**
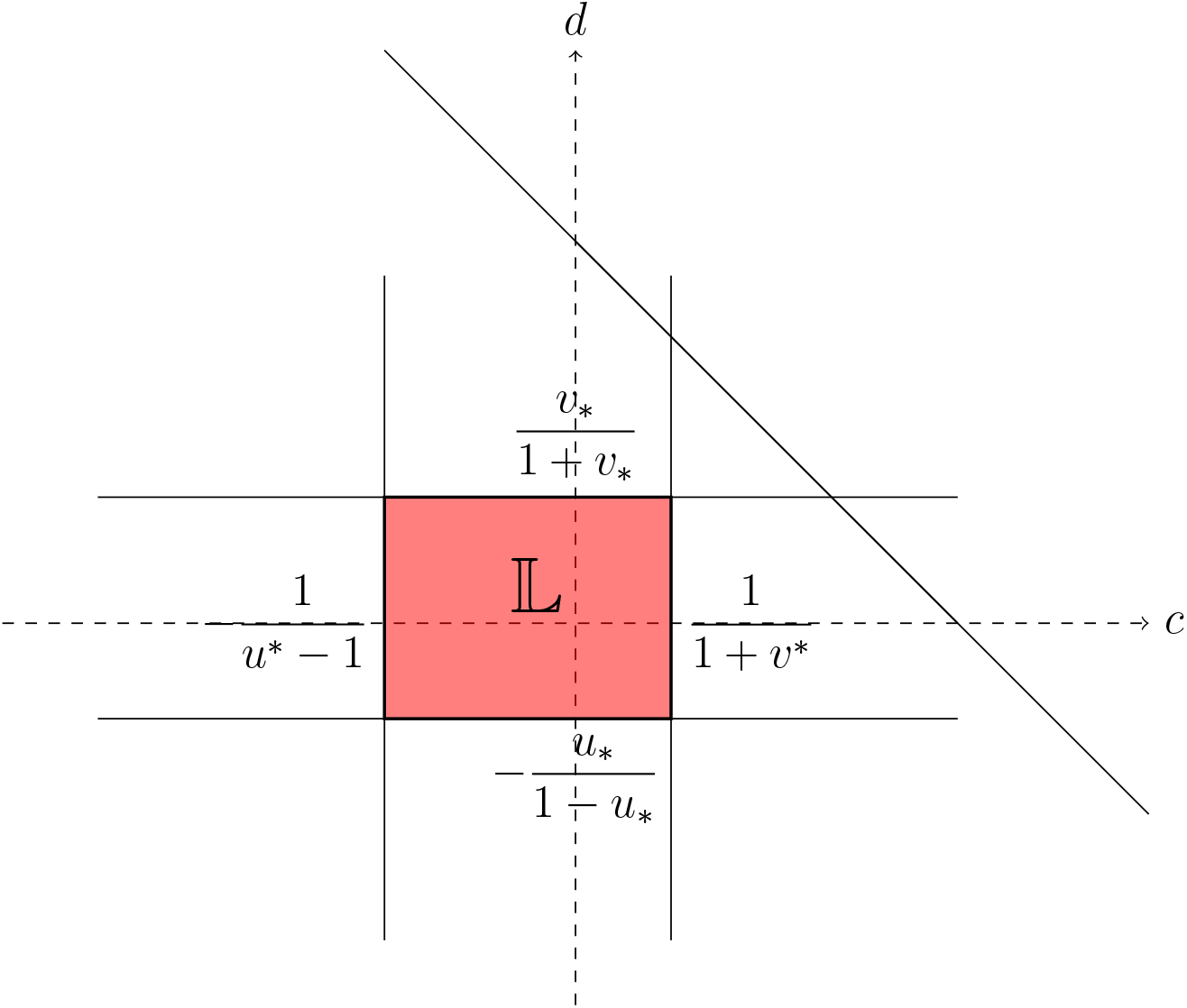
The set of (*c, d*), such that 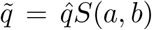 and 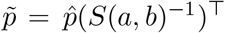 satisfies the conditions (2.1), is given by 𝕃.

